# DiGi: An Integer Programming Formulation for Diploid Genome Inference Using a Pangenome Reference

**DOI:** 10.64898/2026.11.15.766871

**Authors:** Anandita Mishra, Alexander M. Dilthey, Chirag Jain

## Abstract

Compared with genotyping against a single linear reference genome, pangenome-based genotyping enables more comprehensive detection of genetic variation, including structural variants. Existing genotyping methods employ a range of combinatorial, probabilistic, and statistical techniques. One class of methods formulates genotyping as a combinatorial path-finding problem in a pangenome graph, allowing bounded recombinations between reference haplotypes. By restricting solutions to mosaics of reference haplotypes, this formulation implicitly exploits linkage disequilibrium. However, extending this formulation to diploid genomes requires jointly identifying two mosaic paths, leading to an exponential increase in the search space.

To address this challenge, we first represent the pangenome as a *haplotype trellis graph*, which maintains reference haplotypes as separate sequences and permits switching between them only at a predefined set of positions. Second, we formulate a combinatorial optimization problem that uses sequencing reads from a target genome to jointly infer a pair of haplotype paths in the graph. We then develop a probabilistic model that automatically sets the optimization parameters based on the read length and coverage of the input data, and an integer programming formulation to solve the resulting problem exactly. Our implementation, DiGi, successfully reconstructs both paternal and maternal copies of 90% of key HLA genes in target genomes absent from the pangenome reference, achieving sequence identity above 99.9%. On the HG002 Genome in a Bottle benchmark, DiGi achieves favorable F1 scores for genotyping SNVs, indels, and structural variants in the highly polymorphic major histocompatibility complex (MHC) region. DiGi remains accurate across a wide range of sequencing coverages, which makes it well suited to population-scale studies with variable coverage across samples.

**Implementation:** https://github.com/at-cg/DiGi

## 1 Introduction

Population-scale sequencing is reshaping human genetics. UK Biobank, for example, recently completed whole-genome sequencing of half a million individuals [6]. However, the genomic data produced by such initiatives remain heavily skewed towards a few populations. As of October 2026, 88.4% of genome-wide association study participants were of European ancestry. Individuals of continental African ancestry, despite having the greatest genetic diversity of any human population, accounted for only 0.3% [26]. In recent years, national programmes in several low- and middle-income countries (LMICs) have begun to address this imbalance. The GenomeIndia project, for instance, sequenced nearly 10,000 individuals from diverse Indian populations [1]. Scaling such efforts to the population level demands genotyping and phasing methods that are both (i) affordable and (ii) accurate across all classes of variation, from single-nucleotide variants to complex structural variants. Pangenome-based genome inference using low-coverage short-read sequencing data offers one promising route.

Following prior work [9,12], we use *pangenome-based genome inference* to refer to the task of determining (i) which alleles represented in the pangenome an individual carries at each position and (ii) where possible, which parental haplotype each allele lies on. Haplotypes recombine during meiosis, and over many generations a newly sampled haplotype resembles a mosaic of haplotypes already present in the population. Li and Stephens [22] formalized this as an approximate copying model where a new haplotype copies segments of sampled ones such that the switches between them reflect historical recombination, and copying errors reflect novel mutations. This model has since been widely used for imputation and phasing [4,5,16] and, more recently, for genome inference using pangenome references [7,8,12,34].

Existing methods for diploid genome inference on pangenome references differ in what they reconstruct, how they search, and how much coverage they need. Alignment-free methods such as PRG [9], KAGE [13] and PanGenie [12] compare the *k*-mers observed in the reads with those expected from the reference. PanGenie, for example, uses a Li–Stephens-style hidden Markov model to infer the pair of reference haplotypes that best explains the allele-specific *k*-mer counts across the bubbles of a pangenome graph. At low coverage, however, these counts become noisy, making it difficult to distinguish whether an allele is present in zero, one, or two copies. The diploid sampling approach in VG [34] applies a similar optimization strategy within 10 kbp blocks, but selects a pair of haplotypes independently in each block, ignoring long-range linkage disequilibrium. Its software documentation recommends a sequencing coverage of at least 20 × to reliably distinguish heterozygous from homozygous *k*-mers.

PHI [8] formulates haploid genome inference as a global combinatorial optimization problem. It finds the path through the pangenome graph that best agrees with the reads while constraining the number of recombinations. Solved by integer programming, it performs well even at low coverage and reports a full haplotype sequence. Extending its integer programming formulation to diploid genomes is, however, challenging because jointly optimising two paths over a pangenome graph quickly becomes intractable. DipGenie [7] illustrates this trade-off: it reconstructs two haplotypes under a recombination budget, but because its exact integer program does not scale, it falls back on a heuristic. Other methods focus on predefined, short genomic intervals such as specific genes, selecting the best-matching pair of reference haplotypes without allowing recombination [2,30].

Bradbury *et al*. [3] extended genome inference to plant species, whose genomes are often so diverse that much of the sequence cannot be aligned between individuals [35]. They proposed a trellis graph based representation where each reference haplotype defines a horizontal path, and all-vs-all transitions between haplotypes are permitted at orthologous gene boundaries. This design is motivated by high divergence of intergenic regions in plant genomes and the tendency of recombination to occur in regions near genes. Even though their implementation is optimized for plant breeding application, the idea of allowing switches at a sparse set of positions can also help narrow the combinatorial search space for genome inference. Our approach builds on this idea. In this paper, we develop a framework that takes short-read sequencing data from a target genome as input and reconstructs its two haplotype sequences as mosaics of reference haplotypes. Specifically, we make the following contributions:

1. We formalize a *haplotype trellis graph*, in which switches between haplotypes are permitted only at approximately evenly spaced loci that are conserved across individuals. This restriction substantially reduces the search space for diploid genome inference.
2. We formulate diploid genome inference as a combinatorial optimization problem: find two paths through the haplotype trellis graph, with a bounded number of recombinations, whose combined *k*-mer content best matches the reads. The objective explicitly penalizes (i) *k*-mers observed in the reads but absent from the inferred genome, and (ii) *k*-mers in the inferred genome that are absent from the reads. Unlike approaches that require classifying read *k*-mers as homozygous or heterozygous, our formulation avoids this distinction and remains applicable at low sequencing coverage.
3. We prove that this problem is NP-hard and give an integer quadratic programming (IQP) formulation that solves it exactly and runs efficiently in practice. To adapt the method to different read lengths and coverages, we further introduce a probabilistic model that automatically sets the penalty weights in the objective.

We evaluate DiGi on the highly polymorphic human major histocompatibility complex (MHC), for which several complete, haplotype-resolved assemblies are publicly available [24]. Our evaluation comprises five target genomes that are absent from the pangenome. DiGi performs particularly well at low sequencing coverage. At 1 ×, it achieves the highest F1 score for SNVs, indels, and structural variants on HG002 and reconstructs a median of 91.7% of phased gene sequences across the five individuals at ≥ 99.9% identity. At the same coverage and identity threshold, DiGi achieves gene reconstruction rates of 88%–97%, comparing favorably with DipGenie (81%–93%), Locityper (57%-77%), VG (61%–77%), and COSIGT (72%–82%).

## 2 Notation and problem statement

In this section, we formally define the problem. For a string *x*, let |*x*| denote its length, *x*[*i*] its *i*th character (1 ≤ *i* ≤ |*x*|), and *x*[*i, j*] its substring spanning positions *i* through *j*, inclusive. For strings *x* and *y*, we use *x* ° *y* to denote their concatenation. For a positive integer *n*, let [*n*] denote the set {1, 2, …, *n*}. Let ℋ= {*h*_1_, *h*_2_, …, *h*_|_ℋ_|_} be the set of all reference haplotype sequences in a pangenome. Each reference haplotype sequence is a string over alphabet *Σ* = {*A, T, C, G*}.

While inferring the target diploid genome sequence, we want to allow recombinations (haplotype switches) between the pairs of reference haplotypes at known *junctions*. These junctions can be placed at uniformly spaced intervals (discussed later). Let *r* be the number of junctions in a reference haplotype. We assume that these loci are conserved across all |ℋ| reference haplotypes and occur in the same left-to-right order.

Specifically, let 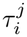 denote the position of *i*th junction in haplotype *h*_*j*_, then we assume that: 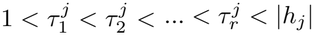 for every 1 ≤ *j* ≤ |ℋ|. We describe how junctions are placed in practice in Section 3.3.

### Haplotype trellis graph

A *haplotype trellis graph* is a directed acyclic graph, representing each haplotype as an horizontal directed chain of vertices, while allowing vertical transitions between haplotypes at fixed junctions (Figure 1a). Formally, it is a directed acyclic vertex-labeled graph *G* = (*V, E, σ*), where *σ* : *V* → *Σ*^∗^ assigns to each vertex, a string over the alphabet *Σ*. We represent each haplotype *h*_*j*_ as an ordered set of vertices 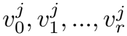 in *G*. These vertices are labeled such that they partition haplotype *h*_*j*_ at *r* junctions, located at positions 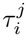 for 1 ≤ *i* ≤ *r*. Specifically, 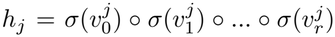, where 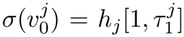, 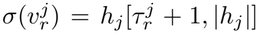, and 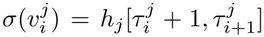 for 0 *< i < r*. One approach to enabling haplotype switching at a junction is to introduce all-to-all connections among the haplotypes (Figure 1a). However, this requires |ℋ|^2^ recombination edges at each junction. Following the optimization in [8], we introduce an empty-labeled junction vertex *w*_*i*_ for each junction *i*∈ [*r*] (Figure 1b). Vertex *w*_*i*_ connects the source and target haplotypes, reducing the number of recombination edges to *O*(|ℋ|) per junction. Finally, we introduce two additional empty-labeled vertices, *s* and *t*, to mark the entry and exit points, respectively, for each path in the graph. In short,

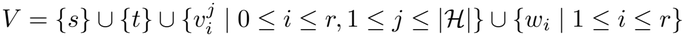

The set of directed edges in *G* is defined as

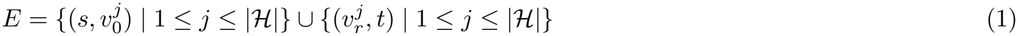

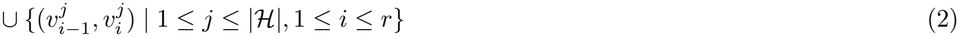

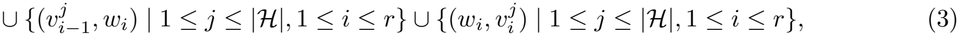

where edges of type (2) join the adjacent substrings of each haplotype and that of type (3) allow a recombination event via junction vertex *w*_*i*_. Previous works [7,8] used the standard pangenome graph representation which merges homologous sequences into shared vertices [15] and they allowed a switch at every vertex in the graph between any pair of haplotypes traversing it. In contrast, our haplotype trellis graph represents haplotypes as disjoint chains and restricts recombination to designated junctions.

**Fig. 1.**
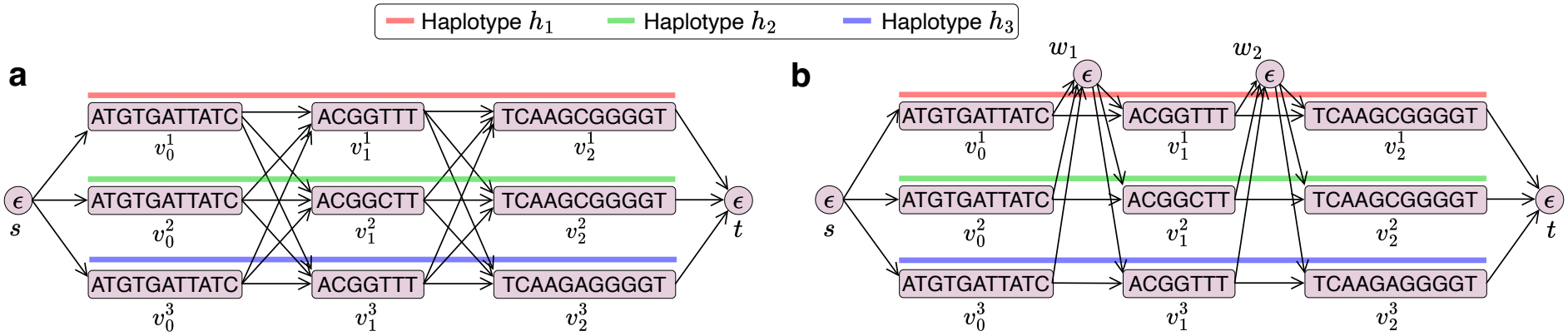
Two equivalent graph representations of three reference haplotypes with two junctions (*r* = 2).

### Inferred path

An *inferred path* through the graph is a path that starts at *s* and ends at *t*. Let *γ*(*P*) denote the number of junction vertices on a path *P*, which equals the number of recombinations along it. Because *P* visits exactly one non-junction vertex in each of the *r* + 1 blocks and one junction per recombination, a path *P* = (*s, u*_1_, …, *u*_*n*_, *t*) has *n* = *r* + 1 + *γ*(*P*) internal vertices. For a string *x*, let *X*_*k*_(*x*) denote the set of distinct *k*-mers in *x*. For an inferred path *P* = (*s, u*_1_, …, *u*_*n*_, *t*), *X*_*k*_(*P*) is the set of distinct *k*-mers in the vertex labels traversed by the path, i.e., 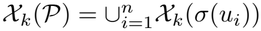. This definition ignores *k*-mers spanning the boundaries between consecutive vertex labels, which in practice represent only a small fraction of all *k*-mers on a path. Let count(*x, y*) denote the number of occurrences of string *y* as a substring of *x*, allowing overlapping occurrences. For an inferred path, 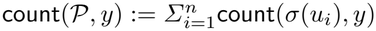, ignoring occurrences spanning vertex-label boundaries.

### Problem statement

Our goal is to infer the target diploid genome as a pair of mosaic paths through the graph. Given sequencing reads from the target genome, we extract the set *S*_*k*_ of distinct *k*-mers observed in the reads. For simplicity of exposition, we assume that all reads are oriented consistently with the target genome. Since the graph contains a large number of candidate *s*-*t* paths, we require an optimization framework to select the most plausible pair of paths. For haploid genome inference, Chandra *et al*. [8] introduced a cost function for path selection that (i) penalizes |*S*_*k*_\*X*_*k*_ (*P*)|, i.e., the number of *missing k-mers* in a path *P*, and (ii) imposes a constant penalty for each recombination. However, a straightforward extension of this objective to diploid genome inference that jointly selects two paths is inadequate. In homozygous regions of the target diploid genome, a single path may account for the observed *k*-mers, leaving the second path unconstrained. To address this issue, the formulation in [7] revised the cost function under the assumption that the *k*-mers extracted from the input reads are pre-partitioned into homozygous and heterozygous sets. We propose a simpler alternative by introducing an additional penalty for occurrences of *novel k-mers*, defined as the *k*-mers in the inferred paths that are absent in *S*_*k*_. For clarity, we first state this objective for a haploid genome and then extend it for a diploid genome. Let *R* denote the maximum number of recombinations in an inferred path.

#### Problem I

*(Haploid path inference problem)*. Determine an inferred path *P* such that *γ*(*P*) ≤ *R* and the following cost is minimized:

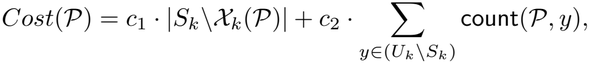

where *U*_*k*_ is the universe set of *k*-mers, and the parameters *c*_1_ and *c*_2_ are non-negative integers representing the penalties for each missing *k*-mer and each occurrence of a novel *k*-mer, respectively.

#### Problem II

*(Diploid path inference problem)*. Determine two inferred paths *P*_1_ and *P*_2_ such that *γ*(*P*_*i*_) ≤*R* for *i* = 1, 2 and the following cost is minimized:

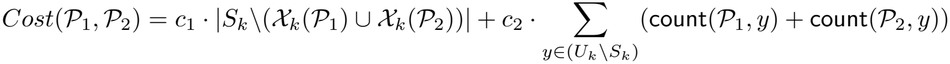

In this paper, we focus on Problem II, motivated by the diploid nature of human genomes. Problem I is known to be NP-hard when the pangenome graph is modeled as general directed acyclic graph with embedded haplotype walks [8]. We show that both Problems I and II remain NP-hard even under the more restrictive haplotype trellis graph representation considered in this paper. Our proof is based on a reduction from the maximum coverage problem and is provided in the Supplementary Document.

#### Theorem 1.

*Problems I and II are NP-hard*.

In our reductions, we set *R* to the number of junction vertices available in the graph. Since an inferred path cannot have more recombinations than that, our proof implies that Problems I and II remain NP-hard even without a bound on *R* (i.e., *R* = ∞).

## 3 Methods

We now present an integer programming formulation for Problem II and a probabilistic model for setting the parameters *c*_1_ and *c*_2_. We then describe the construction of the haplotype trellis graph.

### 3.1 An integer programming solution to Problem II

Recall that *G* = (*V, E, σ*) is a haplotype trellis graph with source *s*, sink *t*, and junction vertices {*w*_*i*_ : *i* ∈ [*r*]}. For a vertex *u* ∈ *V*, let *N* ^+^(*u*) and *N* ^™^(*u*) denote its out- and in-neighbours. Suppose *V*_*k*_ is the set of distinct *k*-mers observed in vertex labels of *G* i.e., 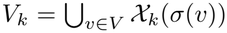. Our goal is to select two *s*–*t* paths *P*_1_ and *P*_2_ that jointly cover the *k*-mers in *S*_*k*_ while containing few *k*-mers outside *S*_*k*_.

#### Decision variables

For *i* ∈ {1, 2}, we define the binary variables

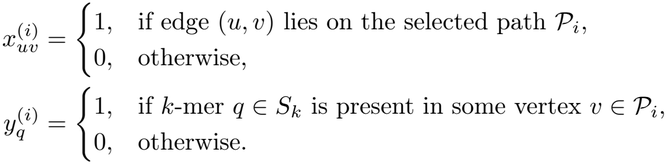

#### Precomputed quantities

We define the edge weight function 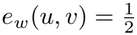 if either *u* or *v* is a junction vertex (so that the cumulative weight of one recombination event is 1) and *e*_*w*_(*u, v*) = 0 otherwise. For each *k*-mer *q* and vertex *v* ∈ *V* \ ({*w*_*i*_ : *i* ∈ [*r*]}∪ {*s, t*}), *m*_*q,v*_ is the number of occurrences of *q* in the label of *v* i.e., *m*_*q,v*_ = count(*σ*(*v*), *q*). *m*_*q,v*_ = 0 for all other nodes.

#### Integer program

We formulate Problem II as the following integer program to jointly identify two paths.

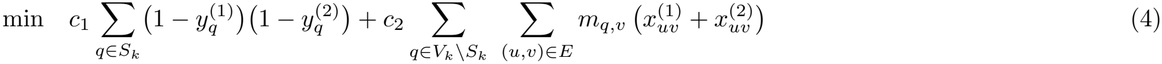

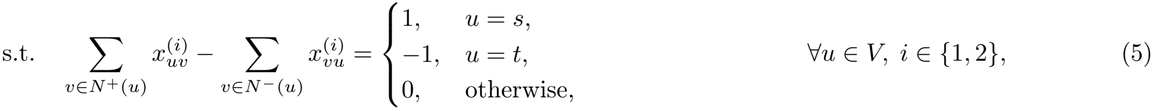

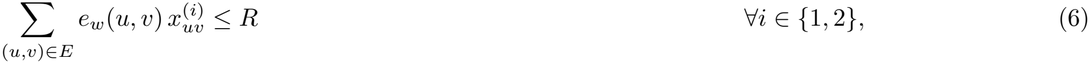

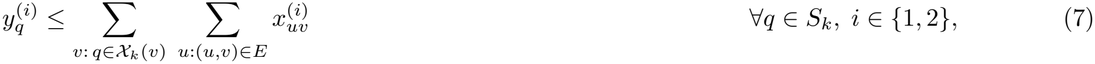

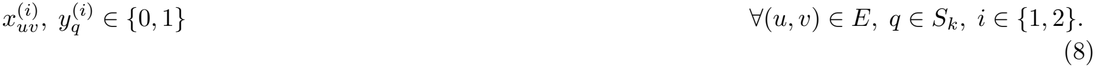

The first term in the objective function (4) counts the *k*-mers of *S*_*k*_ covered by neither path (*missing k-mers*). The second counts occurrences of *k*-mers absent from *S*_*k*_ on the selected paths (*novel k-mers*). Constraint (5) enforces flow conservation, so that each *P*_*i*_ defines an *s*–*t* path. Constraint (6) bounds recombination events on each path according to the user-specified recombination budget *R*. Constraint (7) allows *y*^(*i*)^ = 1 only if *k*-mer *q* occurs in a vertex along *P*_*i*_.

For efficiency, our implementation penalizes missing *k*-mers only among those present in the graph, i.e., we replace *S*_*k*_ with *S*_*k*_ ∩ *V*_*k*_ in the first summation of the objective function (4). This change does not affect the path selection because *k*-mers in *S*_*k*_ that are absent from the graph cannot be covered by any choice of paths, so they add the same constant penalty to every solution. The objective function (4) in our integer program is quadratic in terms of the binary variables 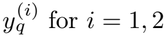 for *i* = 1, 2, whereas all the constraints are linear. The resulting formulation is therefore an integer quadratic program.

### 3.2 Probabilistic model for parameter selection

The parameters *c*_1_ and *c*_2_ are central to our formulation, and their appropriate values depend on sequencing coverage. For example, if the input coverage falls below 1×, then the algorithm must impute missing sequence from the reference haplotypes. The imputed sequence contains *k*-mers not observed in the reads, so the penalty *c*_2_ for novel *k*-mers must be reduced appropriately.

#### Assumptions

We make several simplifying assumptions. First, we assume that the target genome *G*_*T*_ is haploid and represented as a linear sequence. Suppose that mutations in genomes occur only through substitutions. Accordingly, we assume that every sequence spelled by an *s*-*t* path in our haplotype trellis graph has the same length as *G*_*T*_ and differs from *G*_*T*_ only by substitutions. Let *G*_*M*_ denote the sequence spelled by an *s*–*t* path, obtained by concatenating its vertex labels, that minimizes the Hamming distance to *G−* _*T*_. We model the differences between *G*_*T*_ and *G*_*M*_ with a simple independent substitution model, regardless of how the optimization selects *G*_*M*_. At each position, the base is unchanged with probability 1 *µ* and replaced by each of the other three bases with probability *µ/*3.

We assume access to fixed-length sequencing reads of length *L* sampled from *G*_*T*_, with read start positions independently and uniformly distributed along the genome, ignoring boundary effects. Using the Lander-Waterman model [19], the number of reads fully covering a *k*-length interval of *G*_*T*_ is Poisson-distributed with mean 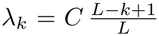, where *C* denotes the sequencing coverage (the total number of sequenced bases divided by the genome length). Next, sequencing errors in a read are assumed to occur independently at each read position. The observed base matches the corresponding base in *G*_*T*_ with probability 1 − *ϵ* and is replaced by each of the other three bases with probability *ϵ/*3.

We also assume that *k* is large enough such that *k*-mers are unique in *G*_*M*_. *k*-mers in *G*_*T*_ that are altered by mutations are assumed not to collide with one another or with *k*-mers in *G*_*M*_. Similarly, sequencing-error-induced *k*-mers are assumed not to collide with *k*-mers in either *G*_*M*_ or *G*_*T*_. We assume that *µ* and *ϵ* are sufficiently small to neglect *k*-mers affected by both mutations and sequencing errors. Thus, each *k*-mer in *S*_*k*_ is assumed to be affected by at most one substitution, arising either from a mutation or from a sequencing error, but not both. We assume that every *k*-mer in each of *S*_*k*_ *\ X* _*k*_(*G*_*M*_) and _*k*_(*G*_*T*_) *\ X* _*k*_(*G*_*M*_) has Hamming distance exactly one from a unique *k*-mer in X_*k*_(*G*_*M*_). Finally, we assume that the presence or absence of each *k*-mer in *S*_*k*_ is independent of that of every other *k*-mer in *S*_*k*_, thereby ignoring dependencies arising from *k*-mers occurring in the same read. *above*,

##### Theorem 2.

*Let p* = Pr(*x* ∈ *S*_*k*_ | *x* ∈ *X*_*k*_(G_*M*_)) *and q* = Pr(*x* ∈ *S*_*k*_ | *x* ∉ *X*_*k*_(G_*M*_)). *Under the assumptions*

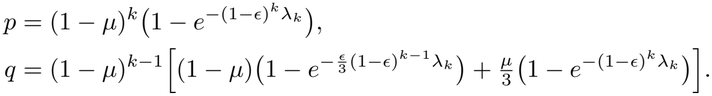

*Setting the parameters of Problem II to*

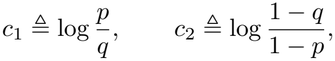

*makes minimizing the cost function in Problem II equivalent to maximizing the log-likelihood of observing the read k-mers S*_*k*_ *given the inferred genome G*_*M*_. *Moreover, p > q (so c*_1_, *c*_2_ *>* 0*) for sufficiently small, positive µ and ϵ*.

*Proof*. Let

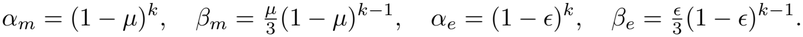

Here *α*_*m*_ and *α*_*e*_ are the probabilities that a *k*-mer in *S*_*k*_ carries no mutation and no error, respectively. *β*_*m*_ and *β*_*e*_ are the probabilities that it carries one specific mutation or error (a given base at a given position). Let *p*_*T*_ and *q*_*T*_ be defined analogously to *p* and *q*, respectively, but with *G*_*T*_ in place of *G*_*M*_. Specifically, *p* = Pr(*x* ∈ *S*_*k*_ | *x* ∈ *X*_*k*_(G_*M*_)) *and qT* = Pr(*x* ∈ *S*_*k*_ | *x* ∉ *X*_*k*_(G_*M*_))

Consider a *k*-mer in *G*_*T*_. The number of reads that cover it and contain it error-free follows a Poisson distribution with mean *α*_*e*_*λ*_*k*_. The number of reads that turn it into a given erroneous *k*-mer follows a Poisson distribution with mean *β*_*e*_*λ*_*k*_. Hence,

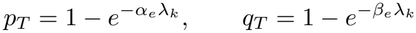

By our assumptions, a *k*-mer of *G*_*M*_ occurs in *G*_*T*_ if and only if it is unaffected by mutations, which occurs with probability *α*_*m*_. In contrast, a *k*-mer in *S*_*k*_ \ *X*_*k*_(*G*_*M*_) may either occur in G_*T*_ if it arises exclusively from a mutation, or be absent from *G*_*T*_ if it arises exclusively from a sequencing error. Therefore

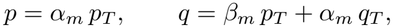

which gives the stated formulas. Moreover, 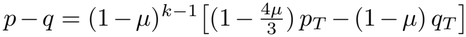. For sufficiently small *µ* and *ϵ*, we have *q*_*T*_ ≪ *p*_*T*_, implying that *p* − *q >* 0. Now the likelihood of observing *S*_*k*_ given G_*M*_ factorizes over the *k*-mers in a fixed universe *U*:

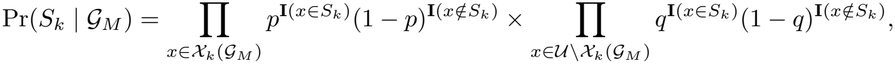

Write *l*_*θ*_(*x*) = **I**(*x* ∈ *S*_*k*_) log *θ* + **I**(*x* ∉ *S*_*k*_) log(1 − *θ*). Then

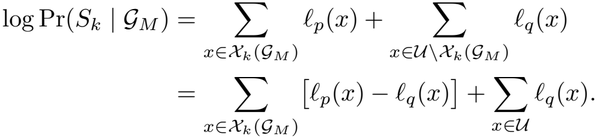

The last sum does not depend on G_*M*_, so

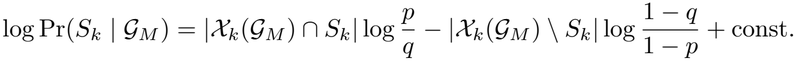

Substitute |*X*_*k*_(*G*_*M*_) ∩ *S*_*k*_| = |*S*_*k*_| − |*S*_*k*_ \ *X*_*k*_(*G*_*M*_)| and absorb the fixed |*S*_*k*_| into the constant. This gives

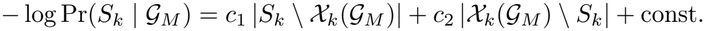

These two counts are the numbers of missing and novel *k*-mers, respectively. Because *k*-mers are assumed to be unique in *G*_*M*_, each novel *k*-mer occurs exactly once. So, counting occurrences, as Problem II does, is the same as counting distinct novel *k*-mers. Hence, with 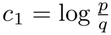 and 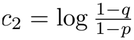, this expression equals the objective of Problem II up to an additive constant.

The above estimation model can be extended to diploid genomes by distinguishing homozygous and heterozygous *k*-mers. For simplicity, we use the haploid parameters in our implementation. We expect this simplification to have little impact on our algorithm, as our formulation considers the presence or absence of each *k*-mer ∈ *S*_*k*_ rather than its copy number in the inferred diploid genome.

### 3.3 Implementation details

We developed a proof-of-concept implementation for genotyping the gene-rich and highly polymorphic major histocompatibility complex (MHC), a 5 Mbp region in the human genome [10].

#### Construction of haplotype trellis graph

We build the pangenome reference as a haplotype trellis graph from two user-supplied inputs: a set of reference haplotype sequences and a set of junction coordinates on GRCh38.

The input MHC haplotype sequences are assumed to be complete, gapless, and on the same strand as the MHC region in GRCh38. Such sequences can be extracted from publicly available haplotype-resolved human genome assemblies [24,25,37]. We place junctions at uniform 10 kbp intervals along GRCh38, which partitions the region into contiguous blocks. Each junction is then lifted over to every reference haplotype. Specifically, we use nf-LO [36] to generate alignment chain files and CrossMap [38] to transfer junction coordinates. We discard any junction that fails to lift over to one or more reference haplotypes. In practice, about 90% of junctions were retained. Finally, we introduce the nodes and edges to the graph as discussed in Section 2.

#### Genome inference

Given short reads from the MHC region of a target genome, we jointly infer its two mosaic paths in the pangenome by solving the IQP formulation (Section 3.1) using Gurobi Optimizer (v13.0.0). We use *k* = 31 throughout our experiments. Read strand orientation is accounted for by representing every *k*-mer in its canonical form. We use FracMinHash sketching [17] with a scaling fraction of 0.05 to subsample *k*-mers and optimize runtime. We use KMC [18] for *k*-mer counting operations. The parameters *c*_1_ and *c*_2_ for the IQP objective are automatically computed in the implementation by using Theorem 2, under the assumption that both the sequencing error rate and the mutation rate equal 0.1% (i.e., *ϵ* = *µ* = 10^™3^). The read length *L* and sequencing coverage *C* are calculated directly from the input data. By default, the maximum number of recombinations is fixed at *R* = 10 per haplotype.

## 4 Results

The inferred haplotype sequences produced by DiGi can serve two downstream applications: (i) genotyping variants by using assembly-based variant callers such as PAV [11], and (ii) recovering phased gene sequences by using annotation tools such as Immuannot [39]. We evaluated DiGi on five human genomes for which both (i) short-read data and (ii) ground-truth MHC haplotypes from long-read assemblies are available.

### Experimental setup

We compared DiGi with PanGenie v4.2.1 [12], DipGenie v1.0 [7], VG v1.76.1 [34], Locityper v1.7.12 [30], and COSIGT v0.2 [2]. For VG, we used its method for computing a personalized pangenome subgraph in diploid haplotype sampling mode [34]. DiGi, DipGenie, VG, and PanGenie each took as input an MHC pangenome and reads sampled from the MHC region of the target genome. DiGi, DipGenie, and VG produce two MHC haplotype sequences, each a mosaic of the reference haplotypes. Pan-Genie produces unphased genotype calls in VCF format. Locityper and COSIGT, by contrast, are targeted genotyping tools that predict phased sequences for user-specified genes, and each predicted sequence is drawn from one of the reference haplotypes in the pangenome. As recommended in their documentation, we supplied these two tools with complete genome assemblies, whole-genome short reads, and the GRCh38 coordinates of the genes of interest. Commands to run each method are available in Supplementary Tables S5 and S6. Each method was run on an Intel Xeon Gold 6248R processor (3.0 GHz, 2×24 cores, 192 GB RAM).

#### Datasets

To construct a pangenome reference for every method, we extracted 108 complete MHC assemblies from publicly available long-read assemblies of 54 individuals [24]. Following standard practice, we aligned each assembly to GRCh38 with minigraph [21] and extracted the sequences aligned to the MHC coordinates (chr6:28,510,125–33,480,577). We then selected five target genomes for genotyping against the pangenome: HG002 (Ashkenazi Jewish), HG005 (Han Chinese), HG01106 (Puerto Rican), HG02055 (African Caribbean), and I002C (Indian). None of these five individuals is among the 54 in the pangenome. Ground-truth MHC haplotype sequences for the five target genomes were obtained from HPRC Release 2 assemblies [25], except for HG002 and I002C, for which we used the T2T assemblies from Hansen *et al*. [14] and Sarashetti *et al*. [32], respectively. Whole-genome short reads are available for all five samples with coverage ranging from 34 to 37×(Supplementary Table S4). We obtained the MHC-specific subset of reads for each sample by aligning its reads to GRCh38. The commands are listed in Supplementary Table S8.

#### Evaluation metrics

We used two ways to assess the accuracy of haplotype sequences computed by DiGi.

- We first evaluated genotyping accuracy in a phase-agnostic manner using the HG002 Genome in a Bottle benchmark v5.0 [29]. We report F1 scores for SNVs, indels, and structural variants (SVs) across a range of short-read coverage depths, from 0.1 × up to full coverage. This comparion includes DiGi, DipGenie, PanGenie, and VG. Note that since every method compared here performs genotyping using a pangenome reference, variants present in the target genome but absent from the pangenome reference haplotypes cannot be recovered by any of them. We excluded Locityper and COSIGT from this experiment because they genotype predefined target loci rather than the entire MHC region.
- Second, we compared each method’s inferred phased sequences of 39 MHC genes with the corresponding ground-truth sequences. We used Immuannot [39] to annotate these 39 genes in the haplotype sequences computed by DiGi, DipGenie, and VG, as well as in the long-read assemblies of the five target genomes. We also ran Immuannot on GRCh38 to obtain the gene coordinates that COSIGT and Locityper require as input. As recommended in their documentation, we extended the interval on both sides for any gene shorter than 10 kbp. PanGenie was excluded from this comparison because it outputs an unphased VCF. Next, for each of the 39 genes in a target genome, agreement between predicted and true sequences was measured as the pairwise sequence alignment identity computed with minimap2 [20]. We also recorded the alignment lengths. Because a target genome would usually carry two copies of every gene, we compared both possible pairings of the two predicted sequences with the two true sequences and kept the pairing that yields the higher total identity.

### Results on the HG002 Genome in a Bottle benchmark

Fig. 2 shows F1 scores, and Supplementary Figs. S1 and S2 show the corresponding precision and recall. At 0.1× and 1× coverage, DiGi achieved the highest F1-scores across SNVs, indels, and SVs. At higher coverages, DiGi’s performance was comparable to that of the best-performing method. PanGenie and VG perform poorly at low coverage because neither was designed for low-coverage data. For example, the VG documentation recommends at least 20× coverage to reliably classify *k*-mers as homozygous or heterozygous. Likewise, PanGenie estimates allele copy number at each bubble of the pangenome graph from the counts of *k*-mers unique to that bubble, and these counts become unreliable when coverage is low. Both DiGi and DipGenie [7] solve global optimization problems with a bound on the number of recombinations, which allows them to exploit linkage disequilibrium present in the reference haplotypes, an advantage that is especially valuable at low coverage. We attribute DiGi’s improved accuracy over DipGenie to three differences: (i) DiGi solves its optimization problem exactly rather than relying on a heuristic, (ii) DiGi does not require read *k*-mers to be classified as homozygous or heterozygous, and (iii) the cost parameters in DiGi’s objective are tuned automatically to the read length and coverage.

**Fig. 2.**
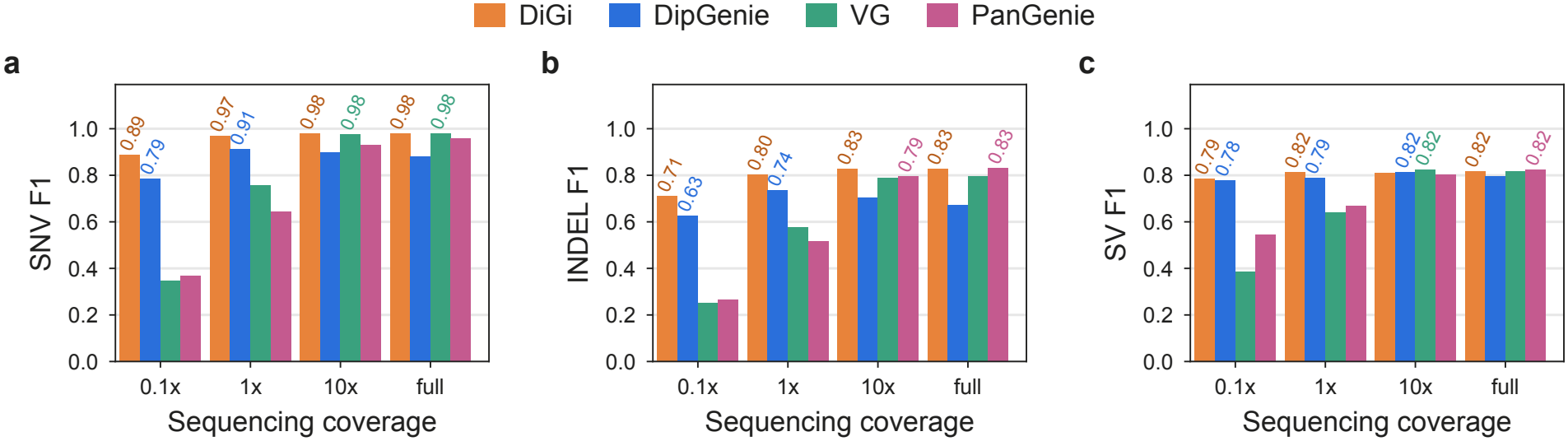
Genotyping accuracy measured using the Genome in a Bottle (GIAB) HG002 assembly-based variant benchmark [29]. F1 scores are shown for sequencing depths ranging from 0.1*×* to full coverage (37*×*). For readability, numerical values are labeled only for the two top-performing methods in each group.

### Haplotype-resolved inference of HLA genes

We evaluated DiGi, DipGenie, VG, Locityper, and COSIGT on their ability to infer phased sequences of 39 MHC genes. Locityper and COSIGT require GRCh38 coordinates for every target gene. Three of the 39 genes (HLA-DRB3, HLA-DRB4, and HLA-Y) are absent from the GRCh38 chromosome 6 primary sequence, so they could not be specified as targets for these two tools. We mark these genes as “not evaluated” for Locityper and COSIGT; all other methods were evaluated on all 39 genes. We ran the comparison at both 1× and full coverage (Fig. 3). At 1 ×, DiGi was the most accurate method, recovering 87.5%–97.2% of gene sequences at ≥ 99.9% identity, compared with 81.2%–92.6% for the next-best method, DipGenie. At full coverage, DiGi’s accuracy increased to 90.3%–98.6%, on par with VG (86.1%–96.9%), Locityper (93.9%–95.7%), and COSIGT (89.4%–95.2%). For Locityper and COSIGT, these percentages are computed over the copies of the 36 genes that could be evaluated, and Fig. 3 shows the remaining three genes as a separate “not evaluated” category. Detailed per-gene alignment statistics for every method are provided in Supplementary Tables S1 and S2.

**Fig. 3.**
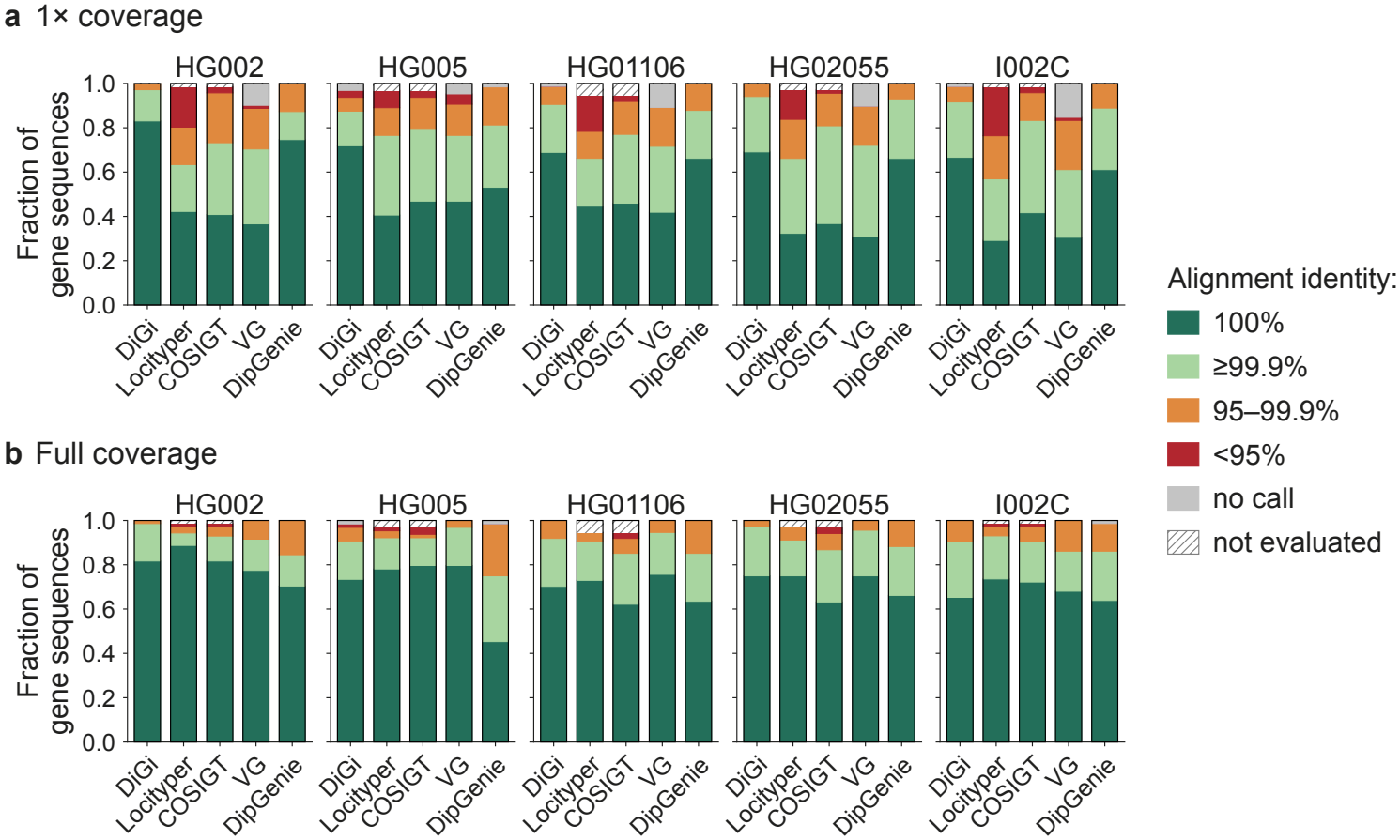
Alignment identity of inferred phased gene sequences for 39 MHC genes across five human genomes, at 1 *×*and full coverage. Most genes are present in two copies per genome, giving 71, 64, 74, 68, and 72 gene copies in total for HG002, HG005, HG01106, HG02055, and I002C, respectively. “not evaluated” marks genes absent from GRCh38 that could not be specified as genotyping targets for Locityper and COSIGT.

### Parameter sensitivity

We next examined how parameter and design choices affect DiGi’s performance. We focused on three: (i) the penalties for missing and novel *k*-mers in inferred paths, controlled by *c*_1_ and *c*_2_ (Section 2), (ii) the limit *R* on the number of recombinations, and (iii) the spacing between junction vertices in the haplotype trellis graph. For each, we compared DiGi with modified versions of itself. We measured accuracy with two metrics commonly used to evaluate genome assemblies: QV (quality value) and *k*-mer completeness [31]. QV is a Phred-scaled estimate of the per-base error rate. QV values of 10, 20, 30, and 40 correspond to one error per 10, 100, 1,000, and 10,000 bp, respectively. It is estimated from the *k*-mers in the inferred sequences that are absent from the ground-truth assembly. *k*-mer completeness is the fraction of *k*-mers in the ground-truth assembly that are also present in the inferred sequences. The two metrics therefore capture different aspects of accuracy.

By default, DiGi sets parameters *c*_1_ and *c*_2_ automatically to non-zero values (Supplementary Table S7). In the optimization objective, *c*_1_ is the cost of each missing *k*-mer, and *c*_2_ is the cost of each occurrence of a novel *k*-mer. To assess their individual contributions, we set each to zero in turn while keeping the other at its default value, and measured the median QV and *k*-mer completeness across the five genomes (Fig. 4a, 4b). The results show that both penalties are needed. With *c*_1_ = 0, inferred paths have no incentive to cover all read *k*-mers, hence *k*-mer completeness drops. With *c*_2_ = 0, paths still include read *k*-mers but nothing prevents them from also including erroneous ones, which lowers QV. This effect is smaller at low coverage where the parameter estimation model already relaxes *c*_2_ so that inferred paths can use haplotype context to include *k*-mers absent from the reads.

**Fig. 4.**
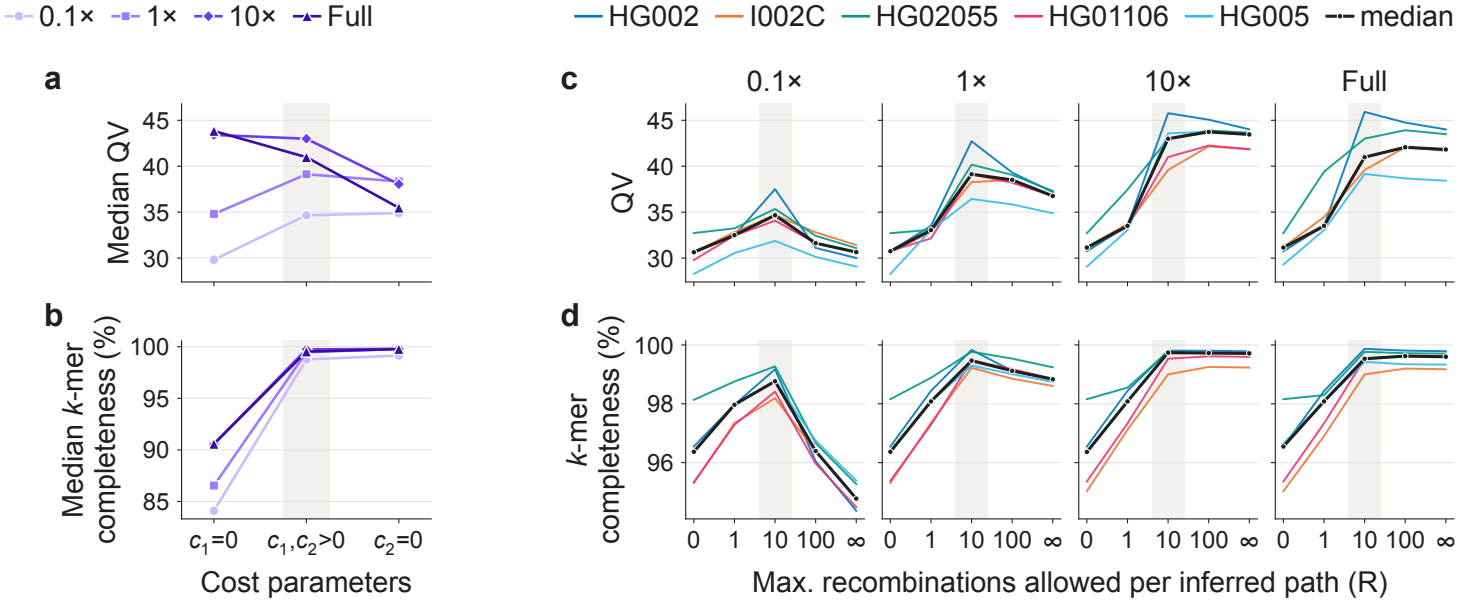
Effect of the optimization parameters *c*_1_, *c*_2_, and *R* on the genome inference accuracy using DiGi. Grey highlighting indicates the default settings.

Unlike *c*_1_ and *c*_2_, which are set automatically, *R* is specified by the user. We used *R* = 10 as default because it gave the best QV and *k*-mer completeness (Fig. 4c, 4d). Setting *R* = 0 reduces inference to selecting the best-matching pair of haplotypes from the pangenome. This is unsuitable for the full MHC of length 5 Mbp, because a new sample is unlikely to match any single reference haplotype along its entire length. At the same time, recombinations must be limited, particularly at low coverage, where sparse read evidence would otherwise let the model switch freely between haplotypes and produce spurious mosaics. We also varied the spacing between junctions in the haplotype trellis graph (Supplementary Fig. S3, S4). Reducing the spacing from 10 kbp to 1 kbp substantially increased runtime because the combinatorial search space for inferred paths grows with the number of junctions, and it also did not improve accuracy. Increasing the spacing to 100 kbp had the opposite effect: runtime decreased, but accuracy dropped.

### Runtime and memory-usage

DiGi completed processing within an hour for every sample and coverage depth, with most of the time spent solving the integer program using Gurobi (Fig. 5). When compared to existing methods, DiGi is faster than DipGenie, comparable to Locityper, and slower than VG, PanGenie, and COSIGT. Runtime and memory usage for all tools are summarized in Supplementary Table S3. DiGi currently runs on a single thread, whereas the other tools used multithreading. For supporting whole-genome genotyping in future, parallel execution should be a straightforward engineering task: each chromosome can be modeled as a separate haplotype trellis graph and solved as an independent integer program.

**Fig. 5.**
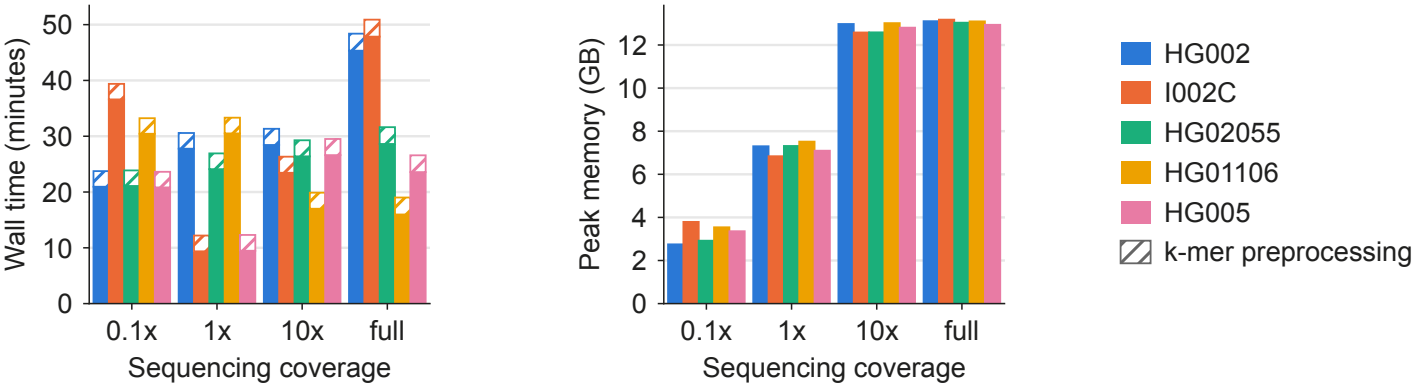
Runtime and peak memory usage of DiGi across the five target genomes and sequencing depths.

## 5 Conclusions and future work

We gave a rigorous formulation for inferring diploid genomes from short reads using a pangenome reference. The haplotype trellis graph keeps reference haplotypes intact and allows switching only at predefined junctions. This design lets DiGi solve the joint inference of two mosaic paths exactly as an integer quadratic program. We also presented a probabilistic model of read *k*-mers and showed that, for suitable choices of *c*_1_ and *c*_2_, solving the problem is equivalent to maximum-likelihood inference under this model. Unlike heuristic search strategies, DiGi returns a provably optimal solution to its objective, and it requires neither a multiple sequence alignment to construct the pangenome graph nor alignment of reads to the pangenome. In practice, DiGi’s advantage is greatest at low coverage, where restricting solutions to mosaics of reference haplotypes compensates for sparse read evidence.

Our results also open several directions for future work. First, the current objective uses only the presence or absence of *k*-mers and treats them as independent. It therefore discards *k*-mer multiplicity and linkage between *k*-mers that come from the same read or read pair. Second, the haplotype trellis graph representation currently assumes complete, gapless haplotypes that are collinear at junctions. These assumptions hold reasonably well for the MHC, but they will need to be relaxed for genome-wide use, where inversions, segmental duplications, and assembly gaps are common. Third, the recombination budget *R* is user-specified. It could instead be derived from haplotype copying models [22], in which the expected number of switches grows with the genetic length of the region and decreases with the number of reference haplotypes. Lastly, near-T2T human genome assemblies are accumulating rapidly [23,25,27,28,33,37]. Because the size of the integer program grows with the number of reference haplotypes, scaling DiGi to pangenomes of more than 1,000 genomes will require further algorithmic optimizations.

## Acknowledgments

This research is supported in part by funding from the DBT/Wellcome Trust India Alliance (IA/I/23/2/506979). A.M. is supported by a Junior Research Fellowship (JRF) from the University Grants Commission (UGC), Government of India. This research used resources of the National Energy Research Scientific Computing Center (NERSC), a U.S. Department of Energy Office of Science User Facility.

## Notes

### Competing Interest Statement

The authors have declared no competing interest.

