## Supplementary Document for "DiGi: An Integer Programming Formulation for Diploid Genome Inference Using a Pangenome Reference"

#### A Supplementary results

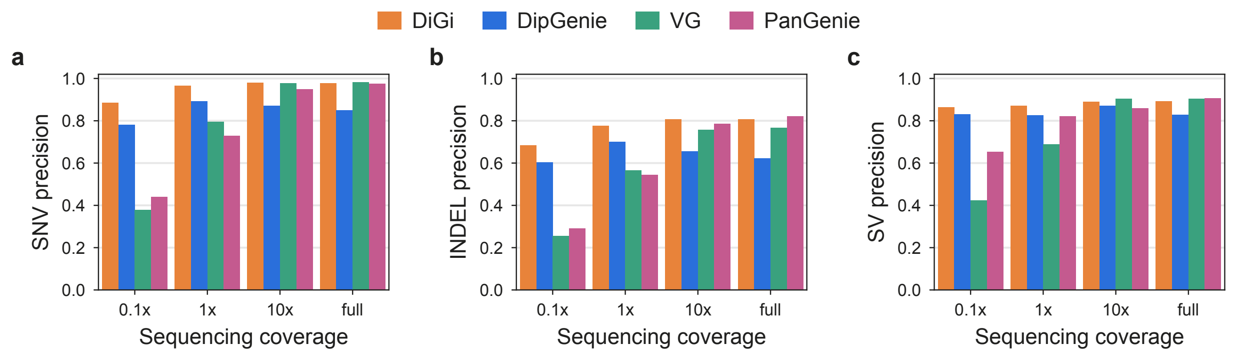

**Fig. S1.** Genotyping accuracy measured using the Genome in a Bottle (GIAB) HG002 variant benchmark. **Precision scores** are shown for sequencing depths ranging from 0.1× to full coverage (37×).

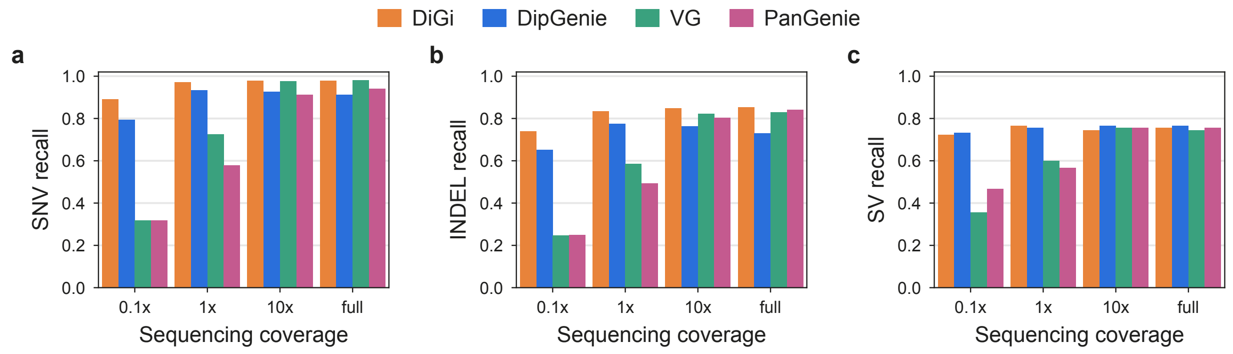

**Fig. S2.** Genotyping accuracy measured using the Genome in a Bottle (GIAB) HG002 variant benchmark. **Recall scores** are shown for sequencing depths ranging from 0.1× to full coverage (37×).

**Table S1.** Per-gene alignment statistics between inferred and true gene sequences at **1× coverage**. Values reported in the table are medians across the five target genomes (HG002, HG005, HG01106, HG02055, and I002C). The columns in the table should be interpreted as following: `Truth`: length of the true gene sequence (bp). `Ratio`: minimap2 alignment length divided by true gene length. `Id`: alignment identity (%). A superscript $c/t$ on the identity value indicates that a tool made $c$ calls for a gene when $t$ were expected in the five target genomes ($c < t$). '×' indicates that the gene was absent from the tool's output. '–' indicates that the gene was not evaluated because it is absent from GRCh38 (`GCA_000001405.15_GRCh38_no_alt_analysis_set.fasta`)

| Gene | Truth | DiGi | | Locityper | | COSIGT | | VG | | DipGenie | |
|---|---|---|---|---|---|---|---|---|---|---|---|
| | | Ratio | Id | Ratio | Id | Ratio | Id | Ratio | Id | Ratio | Id |
| HLA-F | 3544 | 1.000 | 100.00 | 1.000 | 100.00 | 1.000 | 100.00 | 1.000 | 100.00 | 1.000 | 100.00 |
| HLA-V | 1883 | 1.000 | 100.00 | 1.000 | 100.00 | 1.000 | 100.00 | 1.000 | 100.00 | 1.000 | 100.00 |
| HLA-P | 2941 | 1.000 | 100.00 | 1.000 | 100.00 | 1.000 | 100.00 | 1.000 | 100.00 | 1.000 | 100.00 |
| HLA-G | 3138 | 1.000 | 100.00 | 1.000 | 100.00 | 1.000 | 100.00 | 1.000 | 100.00 | 1.000 | 100.00 |
| HLA-H | 3500 | 1.000 | 100.00 | 1.006 | 90.94 | 1.000 | 100.00 | 1.000 | $100.00^{2/7}$ | 1.000 | 100.00 |
| HLA-T | 2486 | 1.000 | 100.00 | 0.681 | 86.98 | 1.000 | 100.00 | 1.000 | $100.00^{2/7}$ | 1.000 | 100.00 |
| HLA-K | 3556 | 1.000 | 100.00 | 1.007 | 84.98 | 1.000 | 100.00 | 1.000 | $99.99^{2/7}$ | 1.000 | 100.00 |
| HLA-U | 731 | 1.000 | 100.00 | 0.568 | 94.23 | 1.000 | 100.00 | 1.000 | $100.00^{2/7}$ | 1.000 | 100.00 |
| HLA-A | 3503 | 1.000 | 100.00 | 1.000 | 99.93 | 1.000 | 100.00 | 0.606 | 97.78 | 1.000 | 100.00 |
| HLA-W | 2950 | 1.000 | 100.00 | 1.001 | 98.46 | 1.000 | 99.98 | 1.000 | 99.05 | 1.000 | 100.00 |
| HLA-J | 3541 | 1.000 | 100.00 | 1.000 | 100.00 | 1.000 | 100.00 | 1.000 | 100.00 | 1.000 | 100.00 |
| HLA-L | 3766 | 1.000 | 100.00 | 1.000 | 99.97 | 1.000 | 99.97 | 1.000 | 99.97 | 1.000 | 99.99 |
| HLA-N | 635 | 1.000 | 100.00 | 1.000 | 100.00 | 1.000 | 99.84 | 1.000 | 99.92 | 1.000 | 100.00 |
| HLA-E | 3822 | 1.000 | 100.00 | 1.000 | 99.97 | 1.000 | 99.97 | 1.000 | 99.97 | 1.000 | 100.00 |
| HLA-C | 4303 | 1.000 | 100.00 | 1.000 | 100.00 | 1.000 | 100.00 | 1.000 | 100.00 | 1.000 | 100.00 |
| HLA-B | 4070 | 1.000 | 100.00 | 1.000 | 99.96 | 1.000 | 99.96 | 1.000 | 99.96 | 1.000 | 99.99 |
| HLA-S | 874 | 1.000 | 100.00 | 1.000 | 100.00 | 1.000 | 100.00 | 1.000 | 100.00 | 1.000 | 100.00 |
| MICA | 12160 | 1.000 | $99.99^{8/10}$ | 0.964 | 85.54 | 1.000 | 99.98 | 1.000 | $100.00^{4/10}$ | 1.000 | 99.99 |
| MICB | 11774 | 1.000 | 99.99 | 1.000 | 99.95 | 1.000 | 99.97 | 1.000 | 99.92 | 1.000 | 99.99 |
| HLA-DRA | 5711 | 1.000 | 100.00 | 1.000 | 100.00 | 1.000 | 99.96 | 1.000 | 99.98 | 1.000 | 100.00 |
| HLA-DRB5 | 12703 | 1.000 | 100.00 | 1.172 | 74.28 | 1.000 | 99.97 | × | $\times^{0/1}$ | 1.000 | 100.00 |
| HLA-DRB1 | 13834 | 1.000 | 99.99 | 0.936 | 87.39 | 0.967 | 97.17 | 0.982 | 98.93 | 1.000 | 99.99 |
| HLA-DQA1 | 6437 | 1.000 | 99.99 | 1.000 | 99.98 | 1.000 | 99.95 | 1.000 | 99.99 | 1.000 | 100.00 |
| HLA-DQB1 | 7131 | 1.000 | 100.00 | 1.000 | 99.98 | 1.000 | 100.00 | 1.000 | 99.96 | 1.000 | 99.99 |
| HLA-DQA2 | 5396 | 1.000 | 100.00 | 1.000 | 99.98 | 1.000 | 99.91 | 1.000 | 100.00 | 1.000 | 99.99 |
| HLA-DQB2 | 7364 | 1.000 | 100.00 | 1.000 | 99.97 | 1.000 | 99.97 | 1.000 | 99.97 | 1.000 | 100.00 |
| HLA-DOB | 4725 | 1.000 | 100.00 | 1.000 | 99.97 | 1.000 | 99.99 | 1.000 | 99.97 | 1.000 | 99.99 |
| TAP2 | 9635 | 1.000 | 99.86 | 1.000 | 99.89 | 1.000 | 99.92 | 1.000 | 99.88 | 1.000 | 99.86 |
| TAP1 | 9270 | 1.000 | 100.00 | 1.000 | 99.96 | 1.000 | 99.99 | 1.000 | 99.98 | 1.000 | 99.99 |
| HLA-DMB | 6712 | 1.000 | 99.99 | 1.000 | 99.97 | 1.000 | 99.96 | 1.000 | 99.98 | 1.000 | 99.94 |
| HLA-DMA | 5011 | 1.000 | 100.00 | 1.000 | 99.98 | 1.000 | 99.95 | 1.000 | 99.98 | 1.000 | 99.97 |
| HLA-DOA | 3653 | 1.000 | 100.00 | 1.000 | 100.00 | 1.000 | 99.99 | 1.000 | 99.99 | 1.000 | 100.00 |
| HLA-DPA1 | 9722 | 1.000 | 100.00 | 1.000 | 99.97 | 1.000 | 99.99 | 1.000 | 100.00 | 1.000 | 100.00 |
| HLA-DPB1 | 11467 | 1.000 | 100.00 | 1.000 | 99.92 | 1.000 | 100.00 | 1.000 | 99.89 | 1.000 | 100.00 |
| HLA-DPA2 | 6743 | 1.000 | 99.99 | 1.000 | 97.35 | 1.000 | 99.95 | 1.000 | 99.94 | 1.000 | 99.99 |
| HLA-DPB2 | 17685 | 1.000 | 99.99 | 1.000 | 99.99 | 1.000 | 99.99 | 1.000 | 99.97 | 1.000 | 99.99 |
| HLA-DRB4 | 14700 | 1.000 | $99.99^{3/4}$ | – | – | – | – | × | $\times^{0/4}$ | 1.000 | $99.99^{3/4}$ |
| HLA-Y | 2254 | 1.000 | $100.00^{2/3}$ | – | – | – | – | 1.000 | $100.00^{1/3}$ | 1.000 | 100.00 |
| HLA-DRB3 | 12806 | 1.000 | 100.00 | – | – | – | – | × | $\times^{0/3}$ | 1.000 | 100.00 |

**Table S2.** Per-gene alignment statistics between inferred and true gene sequences at **full coverage**. Values reported in the table are medians across the five target genomes (HG002, HG005, HG01106, HG02055, and I002C). The columns in the table should be interpreted in the same say as Table S1: `Truth`: length of the true gene sequence (bp). `Ratio`: minimap2 alignment length divided by true gene length. `Id`: alignment identity (%). A superscript $c/t$ on the identity value indicates that a tool made $c$ calls for a gene when $t$ were expected in the five target genomes ($c < t$). '×' indicates that the gene was absent from the tool's output. '–' indicates that the gene was not evaluated because it is absent from GRCh38.

| Gene | Truth | DiGi | | Locityper | | COSIGT | | VG | | DipGenie | |
|---|---|---|---|---|---|---|---|---|---|---|---|
| | | Ratio | Id | Ratio | Id | Ratio | Id | Ratio | Id | Ratio | Id |
| HLA-F | 3544 | 1.000 | 100.00 | 1.000 | 100.00 | 1.000 | 100.00 | 1.000 | 100.00 | 1.000 | 100.00 |
| HLA-V | 1883 | 1.000 | 100.00 | 1.000 | 100.00 | 1.000 | 100.00 | 1.000 | 100.00 | 1.000 | 100.00 |
| HLA-P | 2941 | 1.000 | 100.00 | 1.000 | 100.00 | 1.000 | 100.00 | 1.000 | 100.00 | 1.000 | 100.00 |
| HLA-G | 3138 | 1.000 | 100.00 | 1.000 | 100.00 | 1.000 | 100.00 | 1.000 | 100.00 | 1.000 | 100.00 |
| HLA-H | 3500 | 1.000 | 100.00 | 1.000 | 100.00 | 1.000 | 100.00 | 1.000 | 100.00 | 1.000 | 100.00 |
| HLA-T | 2486 | 1.000 | 100.00 | 1.000 | 100.00 | 1.000 | 100.00 | 1.000 | 100.00 | 1.000 | 100.00 |
| HLA-K | 3556 | 1.000 | 100.00 | 1.000 | 100.00 | 1.000 | 100.00 | 1.000 | 100.00 | 1.000 | 100.00 |
| HLA-U | 731 | 1.000 | 100.00 | 1.000 | 100.00 | 1.000 | 100.00 | 1.000 | 100.00 | 1.000 | 100.00 |
| HLA-A | 3503 | 1.000 | 100.00 | 1.000 | 100.00 | 1.000 | 100.00 | 1.000 | 100.00 | 1.000 | 100.00 |
| HLA-W | 2950 | 1.000 | 100.00 | 1.000 | 100.00 | 1.000 | 100.00 | 1.000 | 100.00 | 1.000 | 100.00 |
| HLA-J | 3541 | 1.000 | 100.00 | 1.000 | 100.00 | 1.000 | 100.00 | 1.000 | 100.00 | 1.000 | 100.00 |
| HLA-L | 3766 | 1.000 | 100.00 | 1.000 | 100.00 | 1.000 | 100.00 | 1.000 | 100.00 | 1.000 | 100.00 |
| HLA-N | 635 | 1.000 | 100.00 | 1.000 | 100.00 | 1.000 | 100.00 | 1.000 | 99.84 | 1.000 | 100.00 |
| HLA-E | 3822 | 1.000 | 100.00 | 1.000 | 100.00 | 1.000 | 100.00 | 1.000 | 100.00 | 1.000 | 100.00 |
| HLA-C | 4303 | 1.000 | 100.00 | 1.000 | 100.00 | 1.000 | 100.00 | 1.000 | 100.00 | 1.000 | 100.00 |
| HLA-B | 4070 | 1.000 | 100.00 | 1.000 | 100.00 | 1.000 | 100.00 | 1.000 | 100.00 | 1.000 | 99.98 |
| HLA-S | 874 | 1.000 | 100.00 | 1.000 | 100.00 | 1.000 | 100.00 | 1.000 | 100.00 | 1.000 | 100.00 |
| MICA | 12160 | 1.000 | 100.00 | 1.000 | 100.00 | 1.000 | 100.00 | 1.000 | 99.99 | 1.000 | 99.99 |
| MICB | 11774 | 1.000 | 99.99 | 1.000 | 99.99 | 1.000 | 99.99 | 1.000 | 99.99 | 1.000 | 99.96 |
| HLA-DRA | 5711 | 1.000 | 100.00 | 1.000 | 100.00 | 1.000 | 100.00 | 1.000 | 100.00 | 1.000 | 100.00 |
| HLA-DRB5 | 12703 | 1.000 | 100.00 | 1.000 | 100.00 | 1.000 | 99.97 | 1.000 | 100.00 | 1.000 | 100.00 |
| HLA-DRB1 | 13834 | 1.000 | 99.99 | 0.961 | 99.74 | 0.967 | 97.17 | 1.000 | 99.98 | 1.000 | $99.92^{9/10}$ |
| HLA-DQA1 | 6437 | 1.000 | 99.99 | 1.000 | 100.00 | 1.000 | 99.95 | 1.000 | 99.99 | 1.000 | 100.00 |
| HLA-DQB1 | 7131 | 1.000 | 100.00 | 1.000 | 100.00 | 1.000 | 100.00 | 1.000 | 100.00 | 1.000 | 100.00 |
| HLA-DQA2 | 5396 | 1.000 | 100.00 | 1.000 | 100.00 | 1.000 | 100.00 | 1.000 | 100.00 | 1.000 | 99.98 |
| HLA-DQB2 | 7364 | 1.000 | 100.00 | 1.000 | 100.00 | 1.000 | 100.00 | 1.000 | 100.00 | 1.000 | 100.00 |
| HLA-DOB | 4725 | 1.000 | 100.00 | 1.000 | 100.00 | 1.000 | 100.00 | 1.000 | 100.00 | 1.000 | 99.99 |
| TAP2 | 9635 | 1.000 | 99.92 | 1.000 | 99.98 | 1.000 | 99.95 | 1.000 | 99.99 | 1.000 | 99.78 |
| TAP1 | 9270 | 1.000 | 100.00 | 1.000 | 100.00 | 1.000 | 100.00 | 1.000 | 100.00 | 1.000 | 99.97 |
| HLA-DMB | 6712 | 1.000 | 99.99 | 1.000 | 100.00 | 1.000 | 99.99 | 1.000 | 100.00 | 1.000 | 99.97 |
| HLA-DMA | 5011 | 1.000 | 100.00 | 1.000 | 100.00 | 1.000 | 100.00 | 1.000 | 100.00 | 1.000 | 99.98 |
| HLA-DOA | 3653 | 1.000 | 100.00 | 1.000 | 100.00 | 1.000 | 100.00 | 1.000 | 100.00 | 1.000 | 100.00 |
| HLA-DPA1 | 9722 | 1.000 | 100.00 | 1.000 | 100.00 | 1.000 | 100.00 | 1.000 | 100.00 | 1.000 | 100.00 |
| HLA-DPB1 | 11467 | 1.000 | 100.00 | 1.000 | 100.00 | 1.000 | 100.00 | 1.000 | 100.00 | 1.000 | 99.97 |
| HLA-DPA2 | 6743 | 1.000 | 99.99 | 1.000 | 100.00 | 1.000 | 100.00 | 1.000 | 100.00 | 1.000 | 99.99 |
| HLA-DPB2 | 17685 | 1.000 | 100.00 | 1.000 | 100.00 | 1.000 | 99.99 | 1.000 | 99.99 | 1.000 | 99.99 |
| HLA-DRB4 | 14700 | 1.000 | $99.99^{3/4}$ | – | – | – | – | 1.000 | 99.99 | 1.000 | $99.97^{3/4}$ |
| HLA-Y | 2254 | 1.000 | 100.00 | – | – | – | – | 1.000 | 100.00 | 1.000 | 100.00 |
| HLA-DRB3 | 12806 | 1.000 | 100.00 | – | – | – | – | 1.000 | 100.00 | 1.000 | 100.00 |

**Table S3.** Wall time and peak memory used by each tool on HG002 short reads, measured with `/usr/bin/time -v`. Time measurements cover the per-sample inference steps only, i.e., building each tool's graph, index or reference panel is not counted, except DiGi's graph $k$-mer generation step (approx. 3 minutes), which is charged to every sample. DiGi does not support multithreading. Hence, DiGi, including the Gurobi solver, used one thread only. All other tools were run with 48 threads. The best values at each coverage are in bold.

| Tool | Threads | 0.1× | 1× | 10× | Full |
|---|---|---|---|---|---|
| DiGi | 1 | 23.7 | 30.6 | 31.3 | 48.4 |
| DipGenie | 48 | 48.5 | 95.9 | 163.6 | 190.8 |
| PanGenie | 48 | **0.9** | **0.8** | **0.8** | **0.9** |
| VG | 48 | 1.3 | 1.4 | 1.4 | 1.5 |
| Locityper† | 48 | –‡ | 6.1 | 13.2 | 52.3 |
| COSIGT† | 48 | 4.2 | 4.1 | 4.9 | 27.8 |

**(a)** Wall time (minutes)

| Tool | 0.1× | 1× | 10× | Full |
|---|---|---|---|---|
| DiGi | 2.8 | 7.3 | 13.0 | 13.1 |
| DipGenie | 41.2 | 66.2 | 108.8 | 112.9 |
| PanGenie | 23.2 | 23.2 | 23.2 | 23.2 |
| VG | 0.7 | 0.6 | 0.7 | 1.4 |
| Locityper† | –‡ | 12.3 | 12.6 | 12.6 |
| COSIGT† | **0.5** | **0.5** | **0.6** | **0.6** |

**(b)** Peak memory (GB)

†Run on whole-genome sequencing reads over 36 MHC genes (excluding HLA-Y, HLA-DRB3, and HLA-DRB4); all other tools were run on MHC reads over 39 genes.

‡Locityper `preproc` failed at 0.1× because its background region contained too few read pairs to estimate the insert-size distribution.

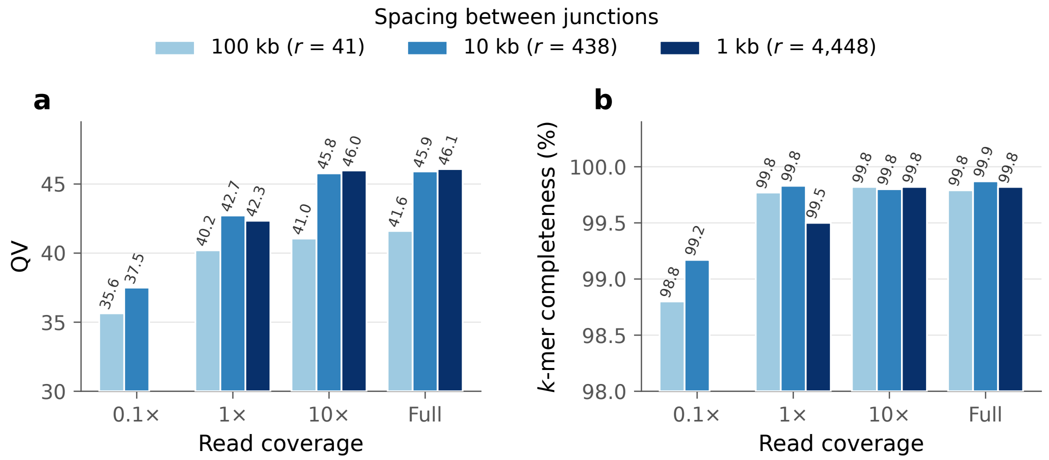

**Fig. S3. Accuracy of haplotypes inferred using DiGi at different junction spacings.** DiGi uses 10 kbp spacing by default, which yields $r = 438$ junctions in the haplotype trellis graph built from 108 MHC haplotypes. At 100 kbp spacing, QV drops, likely because each inferred path must follow a single haplotype over longer stretches. Conversely, too many junctions reduce $k$-mer completeness at low coverage (1×). Results for 0.1× coverage with 1 kbp spacing are omitted because inference exceeded the 48-hour time limit.

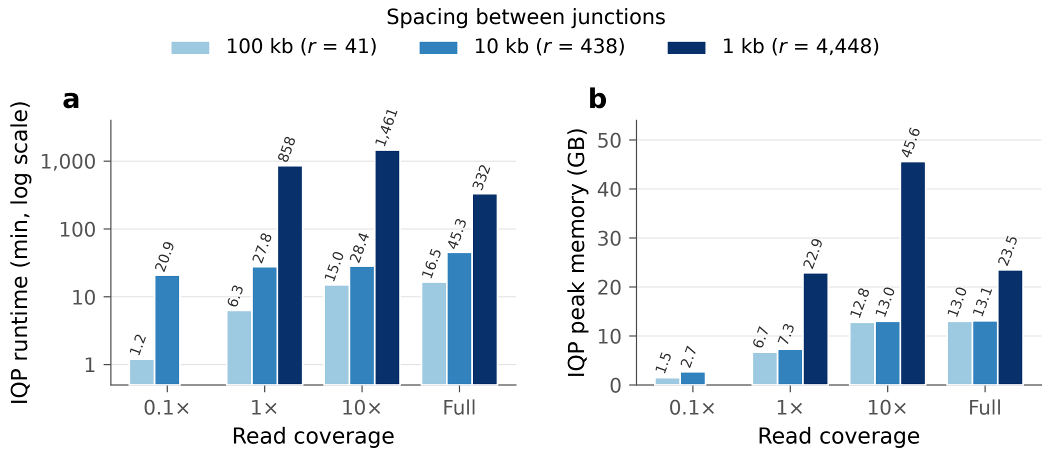

**Fig. S4. Runtime and peak memory usage of DiGi's IQP at different junction spacings.** Runtime and peak memory increase sharply as junctions become denser. Accordingly, moderate junction spacing (∼10 kbp) is essential for keeping the integer program tractable. Results for 0.1× coverage with 1 kbp spacing are omitted because inference exceeded the 48-hour time limit.

### B Data sources and reproducibility

#### B.1 Data sources

**Table S4.** Sources of short-read sequencing data for the five target genomes. Mean whole-genome alignment depth was computed with Samtools against GRCh38 (`GCA_000001405.15_GRCh38_no_alt_analysis_set.fasta`).

| Genome | Sequencing platform | Source | Read length | Mean depth |
|---|---|---|---|---|
| HG002 | HiSeq 2500 | GIAB | 2×250 bp | 34.42× |
| HG005 | NovaSeq 6000 | HPRC R2 | 2×150 bp | 33.97× |
| HG01106 | NovaSeq 6000 | HPRC R2 | 2×150 bp | 35.70× |
| HG02055 | NovaSeq 6000 | HPRC R2 | 2×150 bp | 35.90× |
| I002C | DNBSEQ-T7 | NCBI SRA | 2×150 bp | 36.83× |

**Download URLs.** Reads for HG002 were obtained from the GIAB FTP site (https://ftp-trace.ncbi.nlm.nih.gov/ReferenceSamples/giab/data/AshkenazimTrio/HG002_NA24385_son/NIST_Illumina_2x250bps/reads/). Sequencing data for HG005, HG01106, and HG02055 were retrieved from HPRC Release 2 (`s3://human-pangenomics/working/`), and reads for I002C are available from NCBI SRA under BioProject PRJNA1150503.

#### B.2 Commands for benchmarked genotyping tools

**Table S5. Command lines used to run each genotyping tool.** Each tool is run on 48 threads. Reference construction and preprocessing steps for DipGenie, PanGenie and VG are summarized in Supplementary Table S6.

<table>
<tr><th>Tool</th><th>Command</th></tr>
<tr><td>DipGenie</td><td>./DipGenie -t48 -p2 -R18 -k31 -g <pangenome-MHC-54.gfa.gz> -r <reads.fastq> -o <out></td></tr>
<tr><td>PanGenie</td><td>PanGenie -f <index-prefix> -i <reads.fastq> -s <sample> -j 8 -t 48 -o <out-prefix><br>cat <outfile-prefix>_genotyping.vcf | python3 convert-to-biallelic.py<br><outdir>/multisample-vcfs/callset-filtered.vcf.gz ><br>pangenie_genotyping_biallelic.vcf</td></tr>
<tr><td>VG</td><td>vg haplotypes -v 2 -t 48 -include-reference -diploid-sampling -i <graph>.hapl<br>-k <sample>.MHC.kff -g <sample>.sampled.gbz <graph>.gbz<br>vg paths -x <sample>.sampled.gbz -F -S recombination > <sample>.vg_sampled.fa</td></tr>
<tr><td>Locityper</td><td>locityper preproc -i <R1> <R2> -r <ref.fa> -j <ref>.jf -o <preproc/sample> -@ 48<br>locityper genotype -i <R1> <R2> -d <db> -p <preproc/sample> -o <gt/sample> -@ 48</td></tr>
<tr><td>COSIGT</td><td>make run SMK_ARGS=‘-keep-going’ -cores 48</td></tr>
</table>

**Table S6. Commands used to build each tool's reference panel or index.** DipGenie and VG use the same Minigraph-Cactus graph (first row) but index/preprocess it differently: DipGenie builds its indexes internally as part of its pipeline, while the VG indexing commands are listed here. Locityper builds per-locus haplotype databases directly from the assemblies. PanGenie's graph and index are built from the same haplotypes with its `prepare-graph-from-assemblies` pipeline. COSIGT builds per-locus panels using its main internal Snakemake pipeline mentioned in Table S5. Angle brackets denote placeholders.

| **Tool** | **Command** |
|---|---|
| DipGenie and VG (graph construction) | cactus-pangenome ./js <input_HGSVC_54.txt> -outName <graph> -reference GRCh38 -outDir . |
| VG (index construction) | vg gbwt -G <graph>.gfa -gbz-format -g <graph>.gbz<br>vg index -j <graph>.dist <graph>.gbz<br>vg gbwt -p -num-threads 16 -r <graph>.ri -Z <graph>.gbz<br>vg haplotypes -v 2 -t 16 -H <graph>.hapl <graph>.gbz |
| LociTyper | extract-targets.sh -i <asm_1.fa> -i <asm_2.fa> ... -i <asm_108.fa> -c <loci.bed> -r <ref.fa> -o <haps_dir> -@ <threads><br>locityper target -d <db_dir> -r <ref.fa> -j <ref.jf> -L <haps_dir>/targets.bed -@ <threads><br>locityper align -i <db_dir>/loci/<locus>/haplotypes.fa.gz -o <db_dir>/loci/<locus>/haplotypes.paf.gz -A -@ <threads> |

#### B.3 DiGi Parameter Settings

**Table S7.** Cost parameters $(c_1, c_2)$ computed using Theorem 2 with $\mu = 10^{-3}$, $k = 31$, $\varepsilon = 10^{-3}$, and $R = 10$. $C_{\text{full}}$ denotes the full sequencing coverage over the MHC region, computed as the total number of read bases in the MHC divided by the length of the MHC region in GRCh38. $L$ is the read length. These parameters were used by default for all our experiments.

| | | | 0.1× | | 1× | | 10× | | $C_{full}$ | |
|---|---|---|---|---|---|---|---|---|---|---|
| Sample | $L$ | $C_{full}$ | $c_1$ | $c_2$ | $c_1$ | $c_2$ | $c_1$ | $c_2$ | $c_1$ | $c_2$ |
| HG002 | 250 | 37.3 | 7.3015 | 0.0413 | 7.2041 | 0.4102 | 6.3333 | 3.1184 | 5.1799 | 3.4833 |
| HG005 | 150 | 38.0 | 7.3025 | 0.0375 | 7.2140 | 0.3732 | 6.4045 | 2.9822 | 5.2517 | 3.4836 |
| HG01106 | 150 | 40.0 | 7.3025 | 0.0375 | 7.2140 | 0.3732 | 6.4045 | 2.9822 | 5.2037 | 3.4834 |
| HG02055 | 150 | 40.7 | 7.3025 | 0.0375 | 7.2140 | 0.3732 | 6.4045 | 2.9822 | 5.1875 | 3.4833 |
| I002C | 150 | 37.6 | 7.3025 | 0.0375 | 7.2140 | 0.3732 | 6.4045 | 2.9822 | 5.2616 | 3.4837 |

#### B.4 Commands for other tools

**Table S8.** Other commands used at each step of the analysis.

<table>
<tr><th>Step</th><th>Command</th></tr>
<tr><td>Chain file generation</td><td>nextflow run evotools/nf-LO -source <ref.fa> -target <target_hap.fa><br>-distance near -aligner minimap2 -profile conda -outdir <target_out></td></tr>
<tr><td>Junction liftover</td><td>CrossMap region <target_out>/chainnet/liftover.chain <ref_junctions.bed><br>> <target_junctions> 2> crossmap.log</td></tr>
<tr><td>CRAM to FASTQ</td><td>samtools collate -u -O -reference <cram_ref.fa> <S.cram> S.tmp | samtools<br>fastq -n -s /dev/null -0 /dev/null -1 S_R1.fastq -2 S_R2.fastq</td></tr>
<tr><td>MHC read extraction</td><td>bwa-mem2 mem -R "@RG\tID:S\tSM:S\tPL:PL\tLB:S" <ref.fa> S_R1.fastq.gz<br>S_R2.fastq.gz | samtools sort -o S.bam -<br>samtools view -b -o S.MHC.bam S.bam chr6:28510125-33480577<br>samtools collate -u -O S.MHC.bam S.MHC.tmp | samtools fastq -n -1<br><S.MHC_R1.fastq.gz> -2 <S.MHC_R2.fastq.gz> -s /dev/null -0 /dev/null -</td></tr>
<tr><td>Downsampling reads</td><td>seqtk sample -s100 <S.MHC_R{1,2}.fastq.gz> f > <S.MHC_R{1,2}.Cx.fastq></td></tr>
<tr><td>Gene calling<br>(Immuannot)</td><td>bash scripts.pub.v3/immuannot.sh -c <target.fa> -r Data-2024Feb02<br>-o <target_genes_out></td></tr>
<tr><td>Gene alignment to truth<br>(identity evaluation)</td><td>minimap2 -x asm20 -c -eqx -cs -secondary=no -t 1 <truth_gene.fa><br><inferred_gene.fa></td></tr>
<tr><td>Variant calling on<br>inferred assembly (PAV)</td><td>snakemake -s Snakefile -cores 48 -rerun-incomplete -notemp<br>-restart-times 2 -keep-going</td></tr>
<tr><td>SNV and indel<br>benchmarking</td><td>rtg vcfeval -b HG002_GRCh38_smvar_v50_benchmark_MHC.rebased.vcf.gz<br>-c <exp_smvar.vcf.gz> -t <ref_sdf> -o <out> -output-mode=ga4gh<br>qfy.py <output.vcf.gz> -o <out> -r <ref_fasta><br>-f HG002_GRCh38_smvar_v50_benchmark_MHC_conf.rebased.bed -t ga4gh -V -X</td></tr>
<tr><td>SV benchmarking</td><td>bcftools view -i 'abs(INFO/SVLEN)>50' -Oz -o <exp.SV50.vcf.gz><br><exp.vcf.gz><br>tabix -f -p vcf <exp.SV50.vcf.gz><br>truvari bench -b HG002_GRCh38_stvar_v50_benchmark_MHC.rebased.vcf.gz<br>-c <exp.SV50.vcf.gz><br>-includebed HG002_GRCh38_stvar_v50_benchmark_MHC_conf.rebased.bed<br>-reference GRCh38_no_alt_set_MHC.fasta -o <tru_out><br>truvari refine <tru_out></td></tr>
</table>

#### C NP-hardness Result

**Theorem 1.** *Problems I and II are NP-hard.*

*Proof.* We first prove that Problem I is NP-hard and then extend the reduction to Problem II.
We reduce from the decision version of MAXIMUM COVERAGE PROBLEM: Given a universe $\mathcal{E} = \{e_1, \dots, e_n\}$, a family $\mathcal{F} = \{F_1, \dots, F_m\}$ of subsets of $\mathcal{E}$, and integers $K, L \geq 0$, is there $\mathcal{F}' \subseteq \mathcal{F}$ with $|\mathcal{F}'| \leq K$ and $\left|\bigcup_{F \in \mathcal{F}'} F\right| \geq L$? Without loss of generality, we assume $n \geq 2$, $1 \leq K \leq m$, and no $F_j$ is empty.
*Representation.* Let $b = \lceil \log_3 n \rceil$ and let $\mathrm{ter}(x) \in \{0,1,2\}^b$ be the $b$-digit ternary representation of $x \in [n]$. Define the injective map $\kappa : \mathcal{E} \to \{0,1,2\}^{2b+1}$ by

$$\kappa(e_i) = \mathrm{ter}(i-1) \circ 0^b \circ 1$$

Set $k = 2b + 1$ and $\mathcal{S}_k = \{\kappa(e) \mid e \in \mathcal{E}\}$.
*Haplotypes.* Let $\mathcal{H} = \{h_1, \dots, h_m\}$ and we will construct $h_j$ using $F_j$. Let $\Delta = \max_i |F_i|$. Write $F_j = \{e_{j,1}, e_{j,2}, \dots, e_{j,|F_j|}\}$. Define $e_{j,t} = e_{j,|F_j|}$ for all $|F_j| + 1 \leq t \leq \Delta$. Let

$$\phi_j = \kappa(e_{j,1}) \circ \cdots \circ \kappa(e_{j,\Delta}).$$

$\phi_j$ has fixed length $\ell = k\Delta$, independent of $|F_j|$. Also, every $k$-mer in $\{\kappa(e) \mid e \in F_j\}$ occurs in $\phi_j$. Define $h_j = \phi_j^K$ (i.e., $K$ copies of encoding $\phi_j$), and place $r = K - 1$ junctions at $\tau_i^j = i\ell$.
*Graph.* Let

$$\begin{aligned} V &= \{s,t\} \cup \{v_i^j \mid 0 \leq i \leq K-1,\ 1 \leq j \leq m\} \cup \{w_i \mid 1 \leq i \leq K-1\}, \\ E &= \{(s, v_0^j),\ (v_{K-1}^j, t) \mid 1 \leq j \leq m\} \\ &\quad \cup \{(v_{i-1}^j, v_i^j),\ (v_{i-1}^j, w_i),\ (w_i, v_i^j) \mid 1 \leq i \leq K-1,\ 1 \leq j \leq m\}, \end{aligned}$$

where $\sigma(v_i^j) = \phi_j$. Label $s$, $t$ and all junction vertices $w_i$ with empty strings (see Figure S5).
*Parameters.* Set $c_1 = c_2 = 1$, $R = K - 1$. $G$ has $mK + K + 1$ vertices, $O(mK)$ edges, and labels of length $O(n \log n)$. Since $K \leq m$, the construction is polynomial.

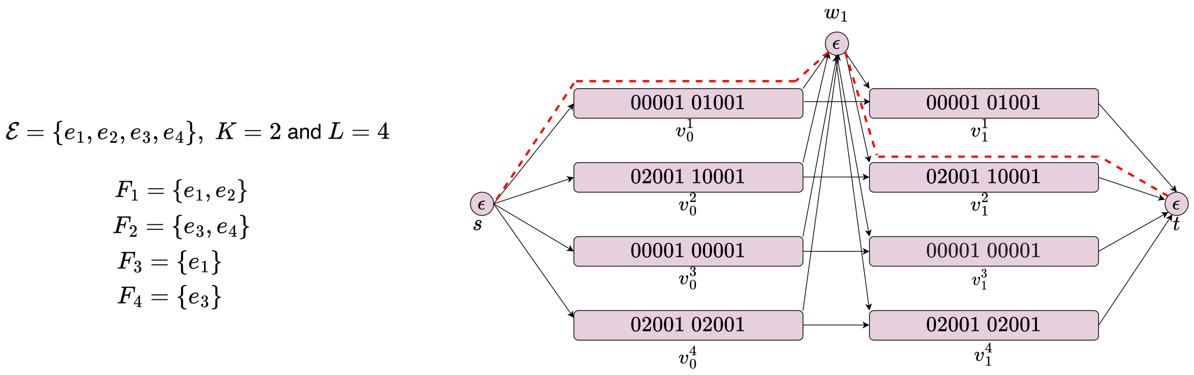

**Fig. S5.** An example of the reduction from an instance of MAXIMUM COVERAGE (left) to the corresponding instance of Problem I (right).

*Cost of a path.* Let $J(\mathcal{P}) \subseteq [m]$ denote the set of distinct haplotype indices covered along an inferred path $\mathcal{P}$. Since $v_{i-1}^j$ has out-edges only to $v_i^j$ and $w_i$, each change of haplotype passes through a junction vertex, so

$$|J(\mathcal{P})| \leq \gamma(\mathcal{P}) + 1. \tag{S1}$$

To evaluate the cost of $\mathcal{P}$, we need to consider the count the occurrences of novel $k$-mers and the count of missing $k$-mers.

(a) *Novel k-mers.* Every element of $\mathcal{S}_k$ has the form $y \circ 0^b \circ 1$, where $y \in \{0,1,2\}^b$. A window of length $k$ in $\phi_j$ starting $o \in \{1, \dots, 2b\}$ characters into an encoding is not in $\mathcal{S}_k$. This is because if $o \leq b$, the encoding's final 1 lies at window position $2b+1-o \in [b+1, 2b]$; if $o > b$, the window ends in the $0^b$ field of the next encoding. Thus only the $\Delta$ aligned windows of $\phi_j$ lie in $\mathcal{S}_k$, and the remaining $(\ell-k+1)-\Delta = 2b(\Delta-1)$ lie in $\mathcal{U}_k \setminus \mathcal{S}_k$. As `count` is summed per vertex, the cost term associated with novel $k$-mers is $C = 2bK(\Delta-1)$ for every $\mathcal{P}$. Accordingly, the cost associated with the number of novel $k$-mers remains fixed irrespective of the choice of $\mathcal{P}$.

(b) *Missing k-mers.* Observe that $\mathcal{X}_k(\mathcal{P}) \cap \mathcal{S}_k = \{\kappa(e) \mid e \in \bigcup_{j\in J(\mathcal{P})} F_j\}$. Hence, by injectivity of $\kappa$, the count of missing $k$-mers is $n - |\bigcup_{j\in J(\mathcal{P})} F_j|$. Accordingly,

$$\mathrm{Cost}(\mathcal{P}) = n - \Big|\bigcup_{j\in J(\mathcal{P})} F_j\Big| + C. \tag{S2}$$

Refer to Figure S5 for one such path (highlighted in red).

*Equivalence.* We next argue that the given instance of Maximum Coverage Problem admits a feasible solution $\mathcal{F}'$ if and only if there exists an inferred path $\mathcal{P}$ in $G$ such that $\mathrm{Cost}(\mathcal{P}) \le \theta$, where $\theta = n - L + 2bK(\Delta-1)$.

($\Rightarrow$) Let $\mathcal{F}' = \{F_{j_1}, \dots, F_{j_p}\}$, $1 \le p \le K$, cover at least $L$ elements. The path that follows $h_{j_1}$, switches to $h_{j_{i+1}}$ at $w_i$ for $1 \le i < p$, and follows $h_{j_p}$ until $t$, has $\gamma(\mathcal{P}) = p-1 \le R$ and $J(\mathcal{P}) = \{j_1, \dots, j_p\}$. From (S2), $\mathrm{Cost}(\mathcal{P}) \le \theta$, where $\theta = n - L + 2bK(\Delta-1)$.

($\Leftarrow$) If $\gamma(\mathcal{P}) \le R$ and $\mathrm{Cost}(\mathcal{P}) \le \theta$, then $\mathcal{F}' = \{F_j \mid j \in J(\mathcal{P})\}$ satisfies $|\mathcal{F}'| \le R+1 = K$ by (S1) and covers at least $L$ elements by (S2).

Hence, Problem I is NP-hard.

*Extension to Problem II.* We reduce from Maximum Coverage Problem with an even budget, i.e., the number of selected sets must be even. This problem remains NP-hard: If an instance $(\mathcal{E}, \mathcal{F}, K', L)$ has odd $K'$, add a set $D$ of $n+1$ fresh elements and pass to $(\mathcal{E} \cup D,\ \mathcal{F} \cup \{D\},\ K'+1,\ L+n+1)$. The sets in $\mathcal{F}$ cover at most $n$ elements, so every solution of the new instance contains $D$, and removing $D$ gives a solution of the original instance. Conversely, adding $D$ to a solution of the original instance gives a solution of the new one.

Now, let the instance be $(\mathcal{E}, \mathcal{F}, 2K, L)$, with the same assumptions as before. Build $G$, $\mathcal{S}_k$, $c_1 = c_2 = 1$ and $R = K-1$ exactly as before. For two inferred paths $\mathcal{P}_1, \mathcal{P}_2$, the cost associated with the occurrences of novel $k$-mers remains $C$ for each of the two paths. The number of missing $k$-mers, i.e., $(\mathcal{X}_k(\mathcal{P}_1) \cup \mathcal{X}_k(\mathcal{P}_2)) \cap \mathcal{S}_k = \{\kappa(e) \mid e \in \bigcup_{j\in J(\mathcal{P}_1)\cup J(\mathcal{P}_2)} F_j\}$. Hence

$$\mathrm{Cost}(\mathcal{P}_1, \mathcal{P}_2) = n - \Big|\bigcup_{j\in J(\mathcal{P}_1)\cup J(\mathcal{P}_2)} F_j\Big| + 2C. \tag{S3}$$

Let $\theta_2 = n - L + 2C$. The given instance $(\mathcal{E}, \mathcal{F}, 2K, L)$ of Maximum Coverage Problem admits a feasible solution $\mathcal{F}'$ if and only if there exist inferred paths $\mathcal{P}_1, \mathcal{P}_2$ in $G$ such that $\mathrm{Cost}(\mathcal{P}_1, \mathcal{P}_2) \le \theta_2$.

($\Rightarrow$) Let $\mathcal{F}' = \{F_{j_1}, \dots, F_{j_p}\}$, $1 \le p \le 2K$, cover at least $L$ elements, and let $q = \min(p, K)$. Build $\mathcal{P}_1$ from the index sequence $(j_1, \dots, j_q)$ and $\mathcal{P}_2$ from $(j_{p-q+1}, \dots, j_p)$, as in the haploid case. The two paths coincide when $p \le K$. Then $\gamma(\mathcal{P}_1) = \gamma(\mathcal{P}_2) = q-1 \le K-1$ and $K-1 = R$. Since $p \le 2q$, $J(\mathcal{P}_1) \cup J(\mathcal{P}_2) = \{j_1, \dots, j_p\}$, so by (S3), $\mathrm{Cost}(\mathcal{P}_1, \mathcal{P}_2) \le \theta_2$.

($\Leftarrow$) Suppose $\gamma(\mathcal{P}_1), \gamma(\mathcal{P}_2) \le R$ and $\mathrm{Cost}(\mathcal{P}_1, \mathcal{P}_2) \le \theta_2$. Let $\mathcal{F}' = \{F_j \mid j \in J(\mathcal{P}_1) \cup J(\mathcal{P}_2)\}$. By (S1), $|\mathcal{F}'| \le |J(\mathcal{P}_1)| + |J(\mathcal{P}_2)| \le 2(R+1) = 2K$, and by (S3), $\mathcal{F}'$ covers at least $L$ elements.

Hence Problem II is NP-hard. □
